# Reconciling Conflicting Models for Global Control of Cell-Cycle Transcription

**DOI:** 10.1101/116798

**Authors:** Chun-Yi Cho, Francis C. Motta, Christina M. Kelliher, Anastasia Deckard, Steven B. Haase

**Affiliations:** Department of Biology, Duke University, Durham, North Carolina 27708; Department of Mathematics, Duke University, Durham, North Carolina 27708

**Author notes:** These authors contributed equally to this work (Chun-Yi Cho, Francis C. Motta). Corresponding author. Mailing address: Department of Biology, Box 91000, Durham, NC 27708-1000.

## Abstract

How the program of periodic cell-cycle transcription is controlled has been debated for several years. Models have ranged from control by a CDK-APC/C oscillator, by a transcription factor (TF) network, or by coupled CDK-APC/C and TF networks. In contrast to current models, a recent study concluded that the cell-cycle transcriptional program is primarily controlled by a CDK-APC/C oscillator with little input from the TF network. This conclusion was largely based on an assumption that substantial drops in transcript levels of network TFs would render them unable to regulate their targets. By combining quantitative modeling and an unbiased analysis of the RNA-seq data, we demonstrate that the data from this recent study are completely consistent with previous reports indicating a critical role of a TF network. Moreover, we report substantial transcript dynamics in cells arrested with intermediate levels of B-cyclins, further supporting the model in which oscillating CDK activity is not required to produce phase-specific transcription.

## INTRODUCTION

A temporal program of cell-cycle transcription is observed across multiple species (Cho et al., 2001; Kelliher et al., 2016; Rustici et al., 2004; Spellman et al., 1998; Whitfield et al., 2002). The cell-cycle transcriptional program is characterized by the phase-specific transcription of a large number of genes (~1000 in budding yeast), which can be organized into clusters based on their timing of expression and regulating transcription factors (reviewed in Haase and Wittenberg, 2014). The entire program is repeated in each new cell cycle, so that each of the genes oscillates in concert with successive cell-cycle progression.

Historically, the cell-cycle transcriptional program was thought to be controlled by a biochemical oscillator based on the antagonistic interactions between cyclin-dependent kinases (CDKs) and the anaphase-promoting complex/cyclosome (APC/C) (Amon et al., 1993; Chen et al., 2004; Cross, 2003; Koch et al., 1996). Because CDKs can trigger many phase-specific cell-cycle events, it was easy to imagine that they could also regulate phase-specific transcription of many genes. In support of this model, CDKs are known to phosphorylate and regulate several transcription factors (TFs) that regulate phase-specific transcription (Amon et al., 1993; Costanzo et al., 2004; de Bruin et al., 2004; Holt et al., 2009; Koch et al., 1996; Landry et al., 2014; Moll et al., 1991; Pic-Taylor et al., 2004; Reynolds et al., 2003; Skotheim et al., 2008; Ubersax et al., 2003).

With the advent of systems-level analyses, it became evident that budding yeast has a highly interconnected network of TFs that can activate/repress each other as well as other cell-cycle genes (Lee et al., 2002; Pramila et al., 2006; Simon et al., 2001). These findings led to the recognition that phase-specific transcription might also arise as an emergent property of a TF network, perhaps operating independently of periodic input from CDKs. Support for this idea came from the finding that a large subset of the cell-cycle transcriptional program continued in cells lacking S-phase and mitotic cyclins, as well as in cells with constitutively high mitotic cyclins (Bristow et al., 2014; Orlando et al., 2008). As cyclins and other CDK regulators are expressed periodically as part of the transcriptional program, the finding that a TF network may be able to produce oscillations opened the door for a model in which CDK oscillations were driven by a TF network oscillator (Simmons Kovacs et al., 2012; 2008).

In aggregate, the studies described above suggested that the CDK-APC/C and the TF network might represent semi-independent oscillatory systems that were coupled by the fact that CDK activities regulate the TFs and the TFs regulate transcription of several CDK regulators. However, a recent study suggested that the cell-cycle transcriptional program was directly driven by a CDK-APC/C oscillator, with little or no autonomous ability of the TF network to operate in the absence of the CDK-APC/C oscillator (Rahi et al., 2016).

Here we show that the data produced by Rahi et al. (2016)are fully compatible with previous work supporting the idea that the CDK-APC/C oscillation is not necessary for driving phase-specific transcription (Bristow et al., 2014; Orlando et al., 2008). An essential difference in our analysis and that of Rahi et al. (2016)concerns the issue of whether cell-cycle transcription that occurs at low transcript abundance can be biologically meaningful. Rahi et al. (2016)made an explicit assumption that if there is a 3-fold drop in the transcript levels of a TF gene in cyclin mutant cells (as compared to wild-type cells), then the remaining TF would be incapable of regulating target genes. In contrast, we recognize that given a complex, interconnected network of transcriptional activators and repressors, low-level expression of a TF in cyclin mutants can function to regulate its target genes. Using a quantitative model of the integrated network, we demonstrate that TFs exhibiting large (10-fold) drops in the amplitude of phase-specific transcription can still generate periodic expression in their target genes. The model easily fit the RNA-seq data of both wild type and cyclin mutant cells from Rahi et al. (2016). Lastly, we present further evidence that a large subset of the cell-cycle transcriptional program can be uncoupled from the oscillation of CDK activity in mitotically arrested cells.

We argue that the cell-cycle transcriptional program emerges from the function of a TF network tightly integrated with CDKs (Bristow et al., 2014; Hillenbrand et al., 2016; Orlando et al., 2008; Simmons Kovacs et al., 2012), rather than from the entrainment of individual TFs by periodic CDK activities produced by an autonomous CDK-APC/C oscillator (Rahi et al., 2016). More broadly, our findings highlight the concept that certain network topologies can produce dynamical functions that are robust to changes in transcript levels, and thus low-level oscillations can be biologically relevant.

## RESULTS

### Models for the control of the cell-cycle transcriptional program

One model for the global control of cell-cycle transcription posits that a post-transcriptional oscillatory CDK-APC/C circuit drives transcriptional oscillations by direct phosphorylation of TFs that control clusters of downstream genes (Figure 1A) (Amon et al., 1993; Chen et al., 2004; Cross, 2003; Koch et al., 1996). This model assumes that the CDK-APC/C network functions as an autonomous biochemical oscillator. While this is certainly true in early embryonic cells, where constitutive input from maternal stores of cyclin RNA is sufficient to drive rapid CDK oscillations (Hara et al., 1980; Murray and Kirschner, 1989), it is not clear that the CDK-APC/C network in somatic cells would similarly produce autonomous oscillations at the relevant (much longer) time scale without periodic input from the transcriptional program.

**Figure 1.**
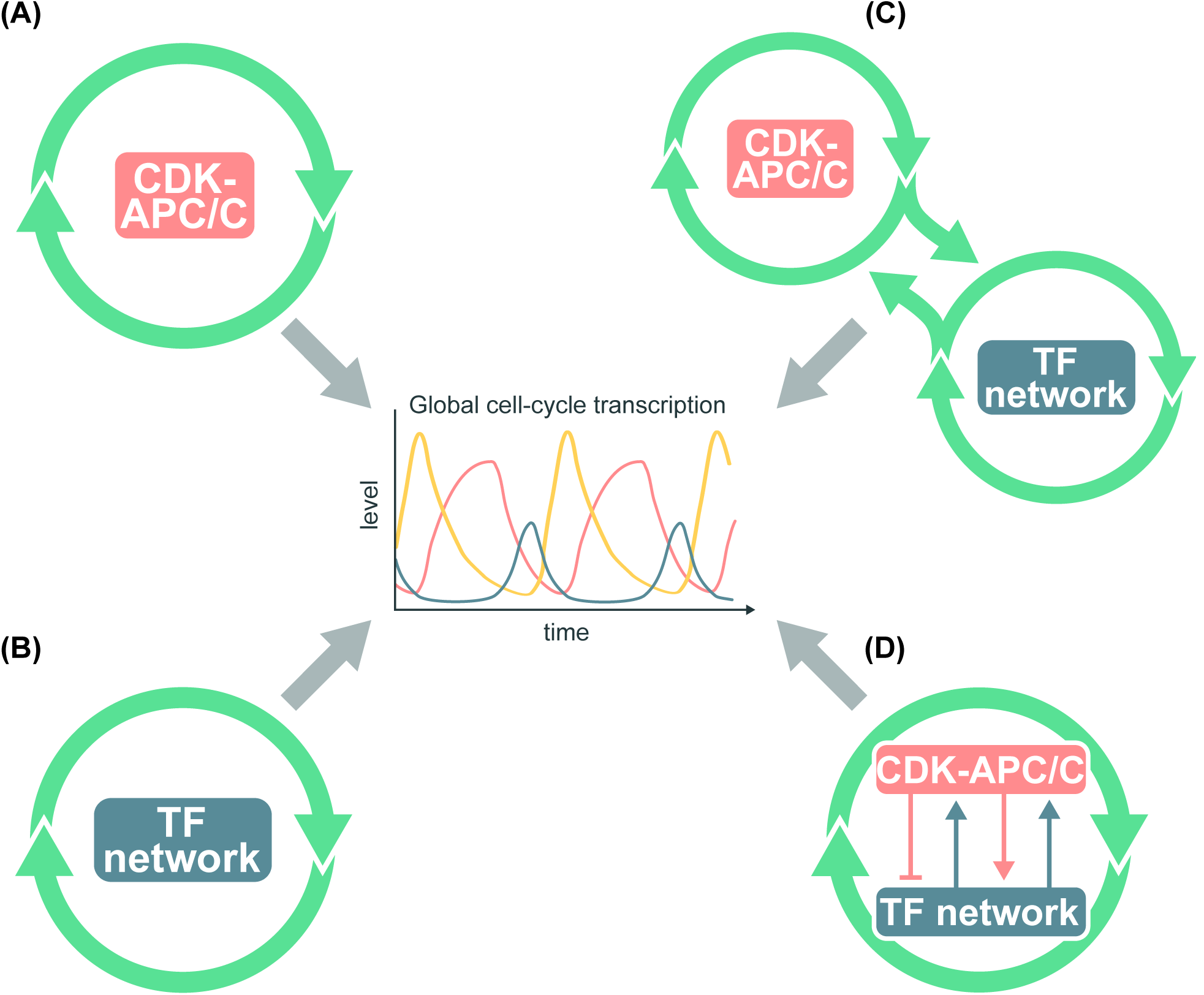
Models for the global control of cell-cycle transcription. (A) The CDK-APC/C network functions as an autonomous oscillator and drives the cell-cycle transcriptional program. (B) The TF network drives the cell-cycle transcriptional program without CDK-APC/C input. (C) The TF network and CDK-APC/C network can function independently, but are coupled to drive the cell-cycle transcriptional program. (D) CDK-APC/C and TF networks are highly connected and act as a single network to control the cell-cycle transcriptional program. Periodic input from CDK-APC/C is not required for oscillations of the transcriptional program.

A second model suggests that phase-specific transcription is brought about by sequential waves of expression of TFs that regulate each other to promote the next wave of expression, with connections between M-phase TFs and G1 TFs restarting the cycle (Figure 1B) (Simon et al., 2001). With appropriate TF activity and stability, such networks could in principle produce phase-specific transcription without input from a CDK-APC/C oscillator. However, it was not clear at the time whether the appropriate TF activity and stability would be obtained without input from CDK-APC/C.

To ask whether phase-specific transcription requires oscillatory input from the CDK-APC/C network, CDK oscillations were blocked either by inhibiting S-phase and mitotic CDK activity or by maintaining a constitutively high level of mitotic CDK activity. Orlando et al. (2008) found that about 70% of phase-specific genes continued to oscillate in cells lacking all S-phase and mitotic cyclins, while cell-cycle events were arrested. Moreover, Bristow et al. (2014) found that many phase-specific genes continued to oscillate in cells depleted of the APC/C coactivator Cdc20, despite the fact that cells were arrested at metaphase. Taken together, these findings suggested that oscillating inputs from the CDK-APC/C network were not required to produce a large subset of phase-specific transcription.

In the experiments by Orlando et al. (2008), about 30% of phase-specific genes were no longer periodically expressed in the mutant cells, suggesting a third model in which the full program of phase-specific transcription requires coupling of the CDK-APC/C network and TF network oscillators (Figure 1C). Subsequent work proposed that the CDK-APC/C oscillator serves as a master oscillator that entrains other autonomous cell-cycle oscillators via a phase-locking mechanism (Lu and Cross, 2010; Oikonomou and Cross, 2010).

When global transcript dynamics were examined in the *cdc28-4/cdk1* cells lacking CDK activities, reproducible transcript oscillations were observed for only a fraction of cell-cycle genes (Simmons Kovacs et al., 2012). Even for those, transcript levels were substantially reduced, and the period of the oscillations was extended. Thus, while CDK oscillations were apparently not critical for phase-specific transcription, some level of CDK activity was required. These findings thus point to a fourth model in which CDK-APC/C and TFs exist in a highly interconnected network (Figure 1D). This model accommodates data from wild-type cells where the entire network oscillates in concert with cell-cycle progression. In various cyclin or APC/C mutants where CDK-APC/C oscillations and cell-cycle progression are halted, the TF network continues to drive oscillations of portions of the cell-cycle transcriptional program.

In a recent publication, Rahi et al. (2016) proposed that the transcriptional oscillations observed in cells lacking S-phase and mitotic cyclins (Orlando et al., 2008) might be related to residual Clb1 left over after the shut-off of *CLB1* expression. They reported that when residual Clb was removed, all phase-specific transcription was halted. Despite this conclusion from transcriptomic data, single-cell analyses indicated that the expression of three genes nevertheless continued to oscillate (Rahi et al., 2016). They concluded that global cell-cycle transcription is predominantly controlled by a CDK-APC/C oscillator, as proposed in early models (Figure 1A). Because this conclusion conflicted with a number of previous findings, we re-examined the data, analyses, and conclusions of this study (Rahi et al., 2016).

### Phase-specific transcription in cells lacking B-cyclins

In the B-cyclin mutant cells (*clb1-6*Δ, denoted as *clb*Δ below) from Orlando et al. (2008), no DNA replication, SPB duplication, mitotic events, or inhibition of bud polarity were observed (Haase et al., 2001; Orlando et al., 2008). Nevertheless, it is possible that there were very low levels of Clb-CDK that were capable of driving the bulk of the transcriptional program while remaining incapable of regulating other cell-cycle events (Rahi et al., 2016). Following a protocol to remove any residual Clb from G1-arrested cells, Rahi et al. (2016)induced a 90- minute pulse of *CLN2* expression and monitored the transcript dynamics in their cyclin mutant cells (*cln1-3*Δ *clb1-*6Δ *MET-CLN2* 0’–90’, denoted as CLN-pulse *clb*Δ below) (Figure 2A). From an analysis of 91 genes, it was concluded that the cell-cycle transcriptional program was severely impaired. Several studies indicated that a much larger number of genes are periodically transcribed in wild-type cells (Figure 2B) (Bristow et al., 2014; Cho et al., 1998; de Lichtenberg et al., 2005; Eser et al., 2014; Granovskaia et al., 2010; Kelliher et al., 2016; Orlando et al., 2008; Pramila et al., 2006; Spellman et al., 1998). To examine the rigor of the conclusion that only three genes continued to oscillate in the *CLN*-pulse *clb*Δ cells, we examined the dynamics of a more comprehensive gene set in the RNA-seq data produced by Rahi et al. (2016) (Figure 2).

**Figure 2.**
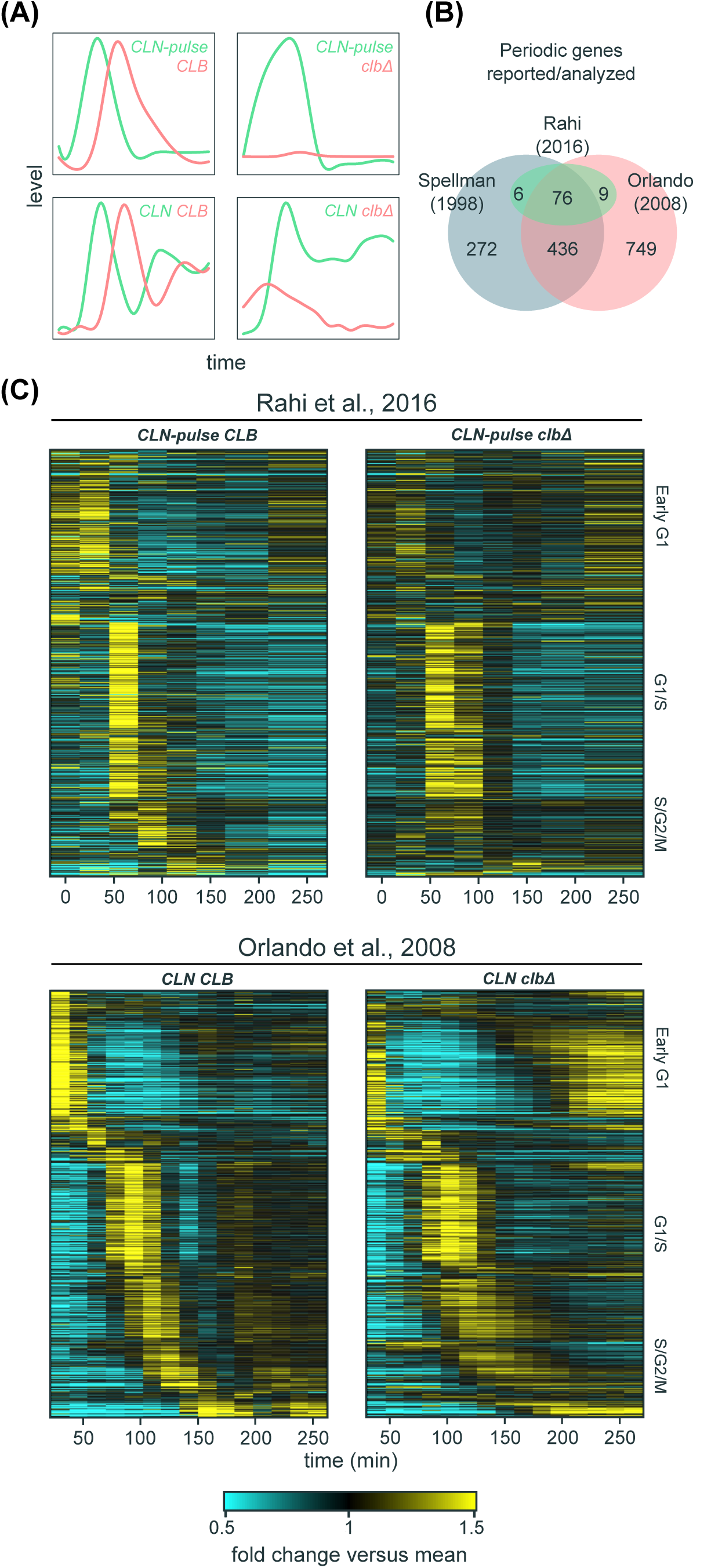
A large program of cell-cycle transcription persists in cells lacking B-cyclins. (A) Cartoon line graphs illustrating the levels of G1 cyclin-CDKs (green) and B-cyclin-CDKs (red) in the time-course experiments from Rahi et al. (2016) (top panels) and Orlando et al. (2008) (bottom panels). (B) Venn diagrams showing the relationships of sets of cell-cycle genes reported previously (Orlando et al., 2008; Spellman et al., 1998) and those examined by Rahi et al. (2016). (C) Heat maps showing transcript dynamics of 881 cell-cycle genes (in the same order) in *CLB* control and *clb*Δ mutant datasets from Orlando et al. (2008) (bottom panels) and Rahi et al. (2016) (top panels). In all experiments, early G1 cells were released into the cell cycle (with the *CLB* expression shut-off for B-cyclin mutants) for time-series gene expression profiling. Transcript levels are depicted as fold change versus mean in individual dataset. Gene lists and corresponding microarray probes can be found in Table S2. See also Figure S1.

In the previous study, Orlando et al. (2008) identified 881 genes (70% of wild-type periodic genes) whose phase-specific transcription remained “on schedule” in the *clb*Δ mutant cells. To ask how these results compared to those reported by Rahi et al. (2016), we directly examined the behaviors of these 881 genes in the RNA-seq data produced by Rahi et al.(2016). In the *CLN-pulse CLB* control (“wild-type-like”), the global dynamics during the first cycle were qualitatively similar to that in the wild-type (*CLN CLB*) cells from Orlando et al. (2008) (Figure 2C, left panels), although distinct dynamical differences appear after the second cycle. Similarly, the transcript dynamics of the first cycle in the *CLN-pulse clb*Δ cells also look qualitatively similar to the *CLN clb*Δ cells from Orlando et al. (2008) (Figure 2C, right panels), including the partial activation of S/G2/M genes. Strikingly, despite the shut-off of *CLN2* expression after 90 minutes in the experiments from Rahi et al. (2016), a second peak of expression for early G1 genes was observed in both *CLN-pulse CLB* control and *CLN-pulse clb*Δ mutant cells (Figure 2C, top panels), reminiscent of the results in the wild type and *clb*Δ mutant cells from Orlando et al. (2008) (Figure 2C, bottom panels).

In both previous studies, the experimental protocols included a media shift at the beginning of the time course (Orlando et al., 2008; Rahi et al., 2016). To ask whether environmental stress response (ESR) (Gasch et al., 2000) or growth rate response (GRR) (Slavov and Botstein, 2011) could be contributing to the transcript dynamics shown in Figure 2C, we eliminated genes that are also in the ESR and GRR program from the 811 genes (Figure S1A). In the remaining 605 genes, similar results were obtained (Figure S1B). Taken together, these analyses demonstrate that a large subset of phase-specific transcription can be produced independently of Clb-CDK activities.

Although global dynamics of all four datasets look similar in the first cycle, clear differences appear in the second cycle. In particular, a second peak of G1/S transcription could be observed in both the wild type and *clb*Δ mutant cells from Orlando et al. (2008) (Figure 2C, bottom panels); however, only one robust cycle of G1/S transcription was observed in the cells from Rahi et al. (2016) (Figure 2C, top panels). These differences likely resulted from the *CLN2* shut-off in the experiments from Rahi et al. (2016) (Figure 2A), as *CLN1/2* are needed for full activation of G1/S transcription partly via inhibition of Whi5 (Skotheim et al., 2008). In the experiments from Orlando et al. (2008), *CLN1/2* were expressed from endogenous promoters and were expressed at relatively high levels (Haase and Reed, 1999; Orlando et al., 2008), likely due to the lack of repression of SBF transcription by Clb2 (Amon et al., 1993; Koch et al., 1996). Presumably, persistent *CLN* expression promoted the re-initiation of G1/S transcription even in the *clb*Δ mutant cells in the experiments from Orlando et al. (2008) (Figure 2C, bottom right). In support of the hypothesis that the shut-off of *CLN2* rather than the depletion of residual Clb impaired the second cycle of transcription, the *CLN-pulse CLB* cells (Figure 2C, top left panel) also have an impaired second cycle of transcription, despite expressing wild-type levels of Clbs. Moreover, it was reported that transcriptional oscillations could persist for some genes if *CLN2* expression was maintained constitutively in the *clb*Δ mutant cells depleted for residual Clb (Rahi et al., 2016).

### Periodicity-ranking algorithms reveal that *SIC1/CDC6/CYK3* are not particularly periodic with respect to other cell-cycle genes in cells lacking B-cyclins

Despite the presence of a large proportion of cell-cycle transcriptional program by visual inspection of the RNA-seq data (Figure 2C), Rahi et al. (2016)reported that only the transcripts of three genes (*SIC1/CDC6/CYK3*) were oscillating in the *CLN*-pulse *clb*Δ cells. This finding was used as a strong argument against the TF network models (Figures 1B and 1C). However, this conclusion was mostly drawn from the analysis of fluorescent microscopic data in single cells, and from an analysis of RNA-seq data restricted to only 91 genes that were grouped together into three clusters. We thus decided to look more globally at periodic transcription within the RNA-seq dataset.

To evaluate the periodicity of all transcripts in the RNA-seq data of the *CLN*-pulse *clb*Δ cells, we utilized two distinct periodicity-ranking algorithms as described and implemented previously (de Lichtenberg et al., 2005; Deckard et al., 2013; Lomb, 1976; Scargle D, 1982) (Materials and Methods). We chose the de Lichtenberg algorithm because Rahi et al. (2016)applied a modified form of this algorithm to conclude that there are no clusters of periodic genes in the *clb*Δ mutant cells. We also utilized the Lomb-Scargle algorithm that was designed to analyze sparsely or unevenly sampled time-series data, such as those produced by Rahi et al. (2016). Due to differences in the sampling density, the periodicity measures returned by the algorithms are not directly comparable between the datasets generated by Rahi et al. (2016) and Orlando et al. (2008). Thus, we took the three genes (*SIC1/CDC6/CYK3*) that Rahi et al. (2016) determined were oscillating and asked where they appeared in the rank-ordered lists produced by the algorithms. Although the two algorithms differ in their quantitative criteria for periodicity (Deckard et al., 2013), both consistently reported that these three genes did not stand out as particularly periodic and were ranked near the bottom of the 91 genes analyzed by Rahi et al. (2016)and at the bottom of the 881 periodic genes shown in Figure 2C (Table 1).

**Table 1.**
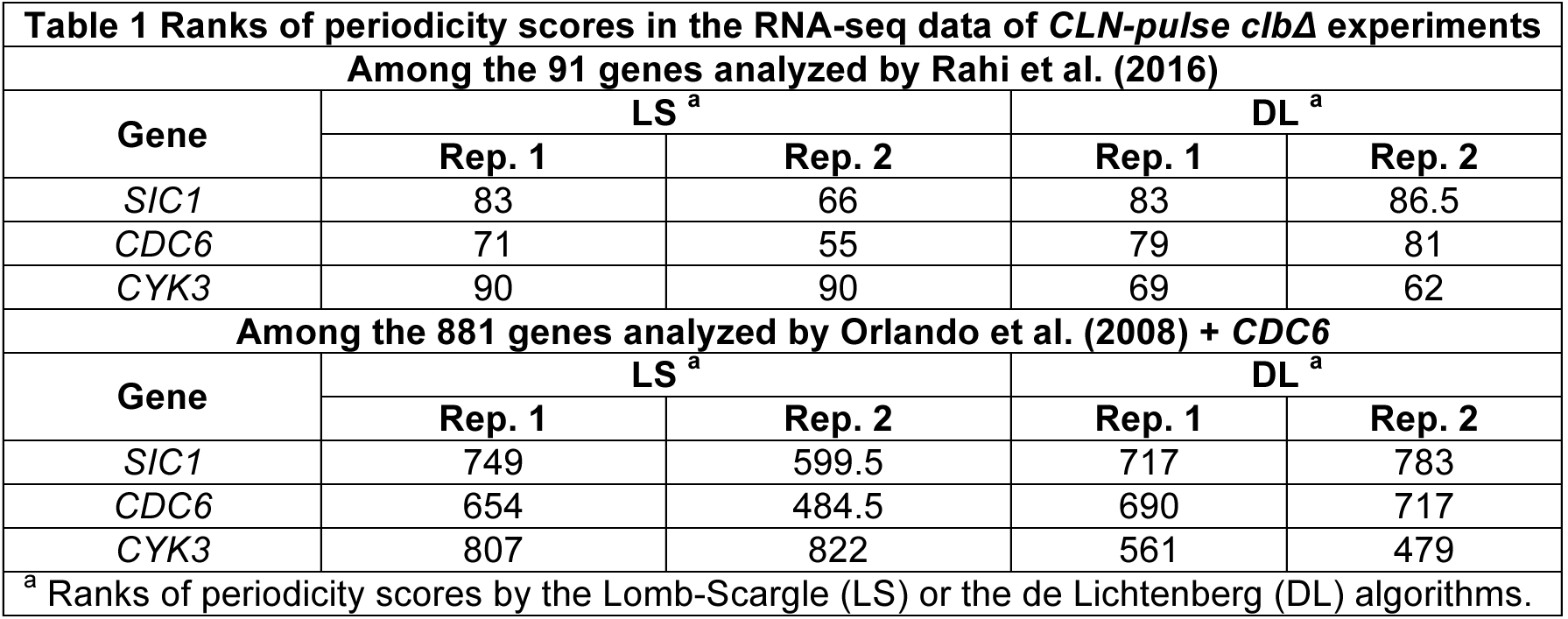
Ranks of periodicity scores in the RNA-seq data of CLN-pulse clbΔ experiments.

|  |  |  |  |  |
| --- | --- | --- | --- | --- |
| **Among the 91 genes analyzed by Rahi et al. (2016)** |  |  |  |  |
| **Gene** | **LS<sup>a</sup>** |  | **DL<sup>a</sup>** |  |
|  | **Rep. 1** | **Rep. 2** | **Rep. 1** | **Rep. 2** |
| *SIC1* | 83 | 66 | 83 | 86.5 |
| *CDC6* | 71 | 55 | 79 | 81 |
| *CYK3* | 90 | 90 | 69 | 62 |
| **Among the 881 genes analyzed by Orlando et al. (2008) + *CDC6*** |  |  |  |  |
| **Gene** | **LS<sup>a</sup>** |  | **DL<sup>a</sup>** |  |
|  | **Rep. 1** | **Rep. 2** | **Rep. 1** | **Rep. 2** |
| *SIC1* | 749 | 599.5 | 717 | 783 |
| *CDC6* | 654 | 484.5 | 690 | 717 |
| *CYK3* | 807 | 822 | 561 | 479 |
| <sup>a</sup> Ranks of periodicity scores by the Lomb-Scargle (LS) or the de Lichtenberg (DL) algorithms. |  |  |  |  |

Whether these three specific genes should be considered periodic in this dataset is certainly arguable. However, it is clear that the conclusion that only three genes oscillated in the B-cyclin mutant cells is not supported by the RNA-seq data, so this data does not argue strongly against the TF network models (Figures 1B and 1C).

### Clusters of phase-specific gene expression are observed in cells lacking B-cyclins

Viewing global transcript dynamics of many genes by heat map can be visually misleading (Figure 2). To look more closely at the dynamics of the cell-cycle transcriptional program in the cyclin mutant cells from Rahi et al. (2016), we examined the behaviors of canonical gene clusters in the budding yeast cell cycle by line graphs (Figure 3) (Haase and Wittenberg, 2014; Spellman et al., 1998). These include the Mcm1 cluster (early G1 genes), SBF/MBF cluster (G1/S genes), histone cluster, Hcm1 cluster (S-phase genes), *CLB2* cluster (G2/M genes), and Swi5/Ace2 cluster (M/G1 genes). For each cluster except the Swi5/Ace2 cluster, two genes whose periodicity was ranked above *SIC1/CDC6/CYK3* in the *CLN-pulse clb*Δ mutant are chosen for illustrations.

**Figure 3.**
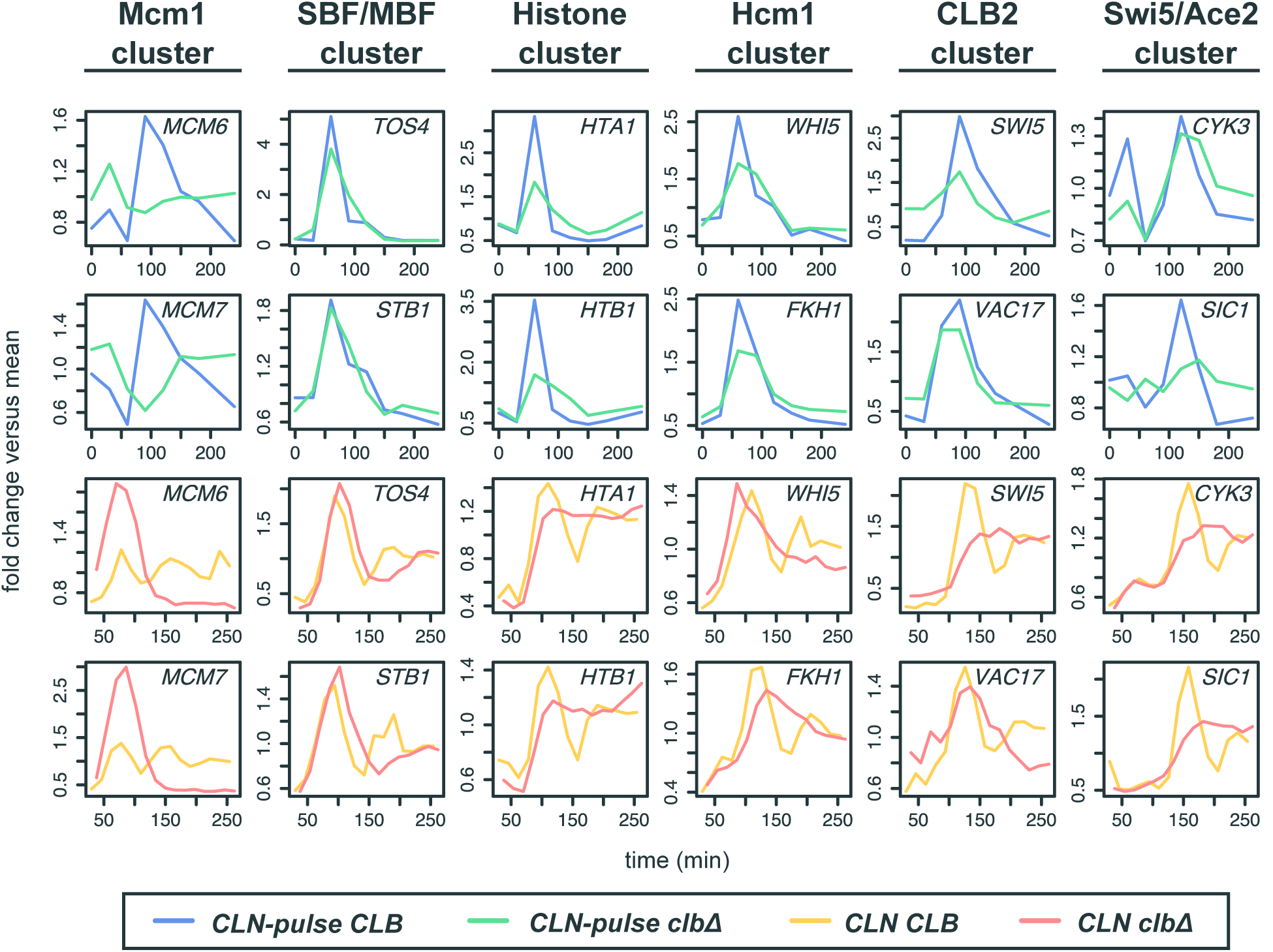
Phase-specific transcription of cell-cycle gene clusters in cells lacking B-cyclins. Line graphs showing transcript dynamics of selected genes in the indicated gene clusters for the *CLB* control and *clb*Δ mutant datasets from Orlando et al. (2008) and Rahi et al. (2016). Early G1 cells were released into the cell cycle (with the *CLB* expression shut-off for B-cyclin mutants) for time-series gene expression profiling. Cln and Clb levels are as shown in Figure 2A. Transcript dynamics are plotted as fold change versus mean as used in Figure 2C. See also Figure S2 and S3.

As shown in Figure 3, the dynamical behaviors (fold change versus mean) of genes from all clusters (except for the Mcm1 cluster) appear remarkably similar in the *CLB* and *clb*Δ strains from the respective studies (Figure 3; blue vs. green lines and orange vs. red lines). These findings strongly suggest that CDK-APC/C oscillations are not required for phase-specific expression and are consistent with the presence of a network of serially activating transcription factors as proposed previously (Figures 1B and 1C) (Orlando et al., 2008; Simmons Kovacs et al., 2012; Simon et al., 2001). The conclusion from Rahi et al. (2016)that the *CLB2* cluster is not activated in the CLN-pulse *clb*Δ cells may have resulted in part from a substantial underrepresentation of the *CLB2* cluster genes in the analysis compared to those previously reported by Spellman et al. (1998). As shown in Figures S2 and S3, among the 30 canonical *CLB2* cluster genes, many were still expressed similarly in both CLN-pulse *CLB* and CLN-pulse *clb*Δ cells. On the other hand, the lack of a strong second pulse for SBF/MBF cluster in the CLN-pulse *clb*Δ cells is consistent with the loss of positive feedback mediated by G1 cyclin-CDKs (Skotheim et al., 2008) as discussed above. Consistently, we observed more prominent transcript dynamics for the canonical clusters in the CLN-pulse *clb*Δ compared to the “CLN-off” *clb*Δ cells that were not induced with *CLN2* expression (Figure S3), suggesting that an input of CDK activity is necessary for the TF network to trigger a cell-cycle transcriptional program with robust amplitude.

### Phase-specific transcription of the network TFs at lower amplitudes in cells lacking 320 B-cyclins

To ask if these behaviors of canonical gene clusters could be regulated in a phase-specific manner by a chain of serially activating TFs as previously proposed (Orlando et al., 2008; Pramila et al., 2006; Simon et al., 2001), we examined the transcript behaviors of the core TFs in the network model (Figure 4A). These genes include *SWI4* (SBF), *HCM1, YHP1, NDD1* (SFF), *SWI5,* and an output *SIC1.* As shown in Figures 4B and 4C, these TF network components exhibited qualitatively similar dynamics with identical temporal order of phase-specific transcription in all four *CLB* control and *clb*Δ mutant datasets (Orlando et al., 2008; Rahi et al., 2016). However, a significant reduction in the amplitude of *SWI5* transcripts was observed in the *CLN-pulse clbΔ* mutant cells, as compared to the CLN-pulse *CLB* cells from Rahi et al. (2016). This observation is consistent with previous studies showing that the activity of SFF is increased by Clb2-CDK, which phosphorylates the components of SFF (Pic-Taylor et al., 2004; Reynolds et al., 2003). Thus, loss of the positive feedback between SFF and Clb2-CDK should decrease the amplitude of SFF targets. A similar but less severe reduction in transcript levels was also observed in the *clb*Δ mutant cells from Orlando et al. (2008) (Figure 4B). It is unclear yet whether the higher level of *SWI5* might have resulted from residual Clb1 or from the fact that *CLN1/2* expression was not shut-off in these experiments.

**Figure 4.**
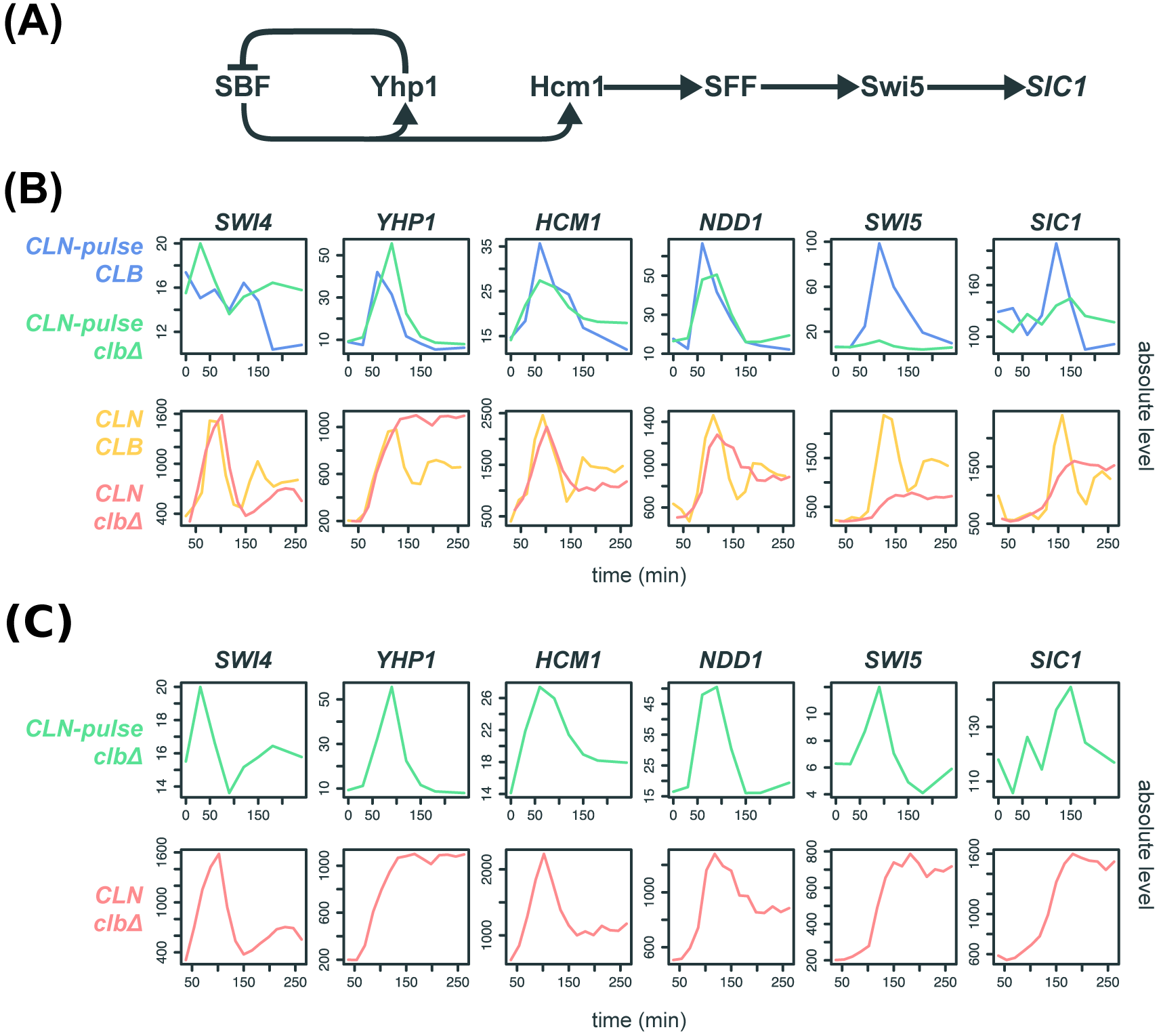
Evidence for serial activation of network TFs in cells lacking B-cyclins. (A) Diagram of the TF network model proposed by Orlando et al. (2008). *SIC1* is an output normally activated by Swi5 during mitotic exit. (B)(C) Line graphs showing the absolute transcript levels (arbitrary units) of the TF network components in the *CLB* control and *clb*Δ mutant datasets from Orlando et al. (2008) and Rahi et al. (2016).

Regardless, the significant drop in the transcript levels for *CLB2* cluster genes (Figure 4B) was highlighted as evidence against the TF network models (Rahi et al., 2016). Paradoxically, Swi5/Ace2 targets, including *SIC1/CDC6/CYK3,* were shown to oscillate in the *CLN-pulse clb*Δ mutant cells without substantial reduction in amplitude (Rahi et al., 2016). Even though the deletion of the *SWI5* gene eliminated the oscillation of *SIC1* transcripts (Rahi et al., 2016), it was concluded that oscillations of *SIC1* transcripts were not controlled by the TF network (Figure 4A) but rather by an undiscovered mechanism.

This conclusion stems from an explicit assumption that a 3-fold reduction in the transcript level of a TF gene compared to wild type would render that TF biologically inactive in terms of the ability to regulate target genes. In the *CLN-pulse clb*Δ mutant cells, *SWI5* and *ACE2* peak at only 8% and 10% of wild-type levels. By applying the “biological significance” test, it was thus concluded that the transcriptional oscillation of *SWI5* could not be driving the transcript dynamics of the *SIC1* gene. Unfortunately, no biological support was given for this assumption. Nonetheless, we wanted to test the assertion that the transcript levels of *SWI5* were too low to produce the transcriptional oscillations of *SIC1* observed in the CLN-pulse *clb*Δ mutant cells.

### A mathematical model demonstrates that the low-amplitude transcriptional oscillations can be biologically relevant in the context of the global network

No explicit quantitative logic was offered for the choice of the biological significance cutoff assumed by Rahi et al. (2016), but it is a common assumption that TFs must achieve some threshold level of expression before they can efficiently activate (or repress) their target genes. This assumption is often made precise with the use of a Hill function nonlinearity in ODE models of transcriptional regulation. Explicitly, if TF A activates transcription of gene B, it is common to model the transcriptional rate of gene B with an ODE model of the form:

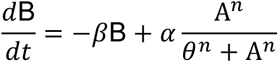
where *θ* represents a (soft) threshold of activation, below which transcription of gene B may be mostly unaffected by levels of TF A, but above which the contribution of TF A to the transcription of gene B is dramatically enhanced.

Using the above equation to consider the activation of the *SIC1* gene by Swi5 in isolation, it is reasonable to assume that the dramatic reduction in the expression of *SWI5* in the *clb*Δ mutant cells would reduce the abundance of Swi5 protein well below the *SIC1* activation threshold. Alternatively, if a very low *SIC1* activation threshold explains the observation of *SIC1* pulses in the *clb*Δ cells, then the equation would predict *SIC1* to be always highly activated in *CLB* cells.

The logic above assumes that there is no additional input to either gene A or B. However, the regulatory interaction between Swi5 and *SIC1* is part of a network with additional inputs including Clb2-CDK and the opposing phosphatase Cdc14 (Figure 5A). The diminished level of *SWI5* transcripts in the *clb*Δ cells is predicted by the network interactions, as Clb2-CDK increases the transcript level of *SWI5* by fully activating SFF. Moreover, Clb2-CDK phosphorylation of Swi5 inhibits its activity by sequestering it in the cytosol (Figure 5A) (Moll et al., 1991; Reynolds et al., 2003). Thus, the timing of *SIC1* activation is not closely tied to the accumulation of Swi5, but rather is delayed until Cdc14 dephosphorylates and activates Swi5 during mitotic exit (Figure 5A) (Visintin et al., 1998). In the *clb*Δ cells, the inhibition of Swi5 by Clb2-CDK is relieved, potentially leading to the robust activation of *SIC1* by low-amplitude expression of *SWI5.* In support of this hypothesis, *SIC1* transcripts accumulated earlier in the *clbΔ* cells than they did in wild-type cells (Figure 4B) (Orlando et al., 2008).

**Figure 5.**
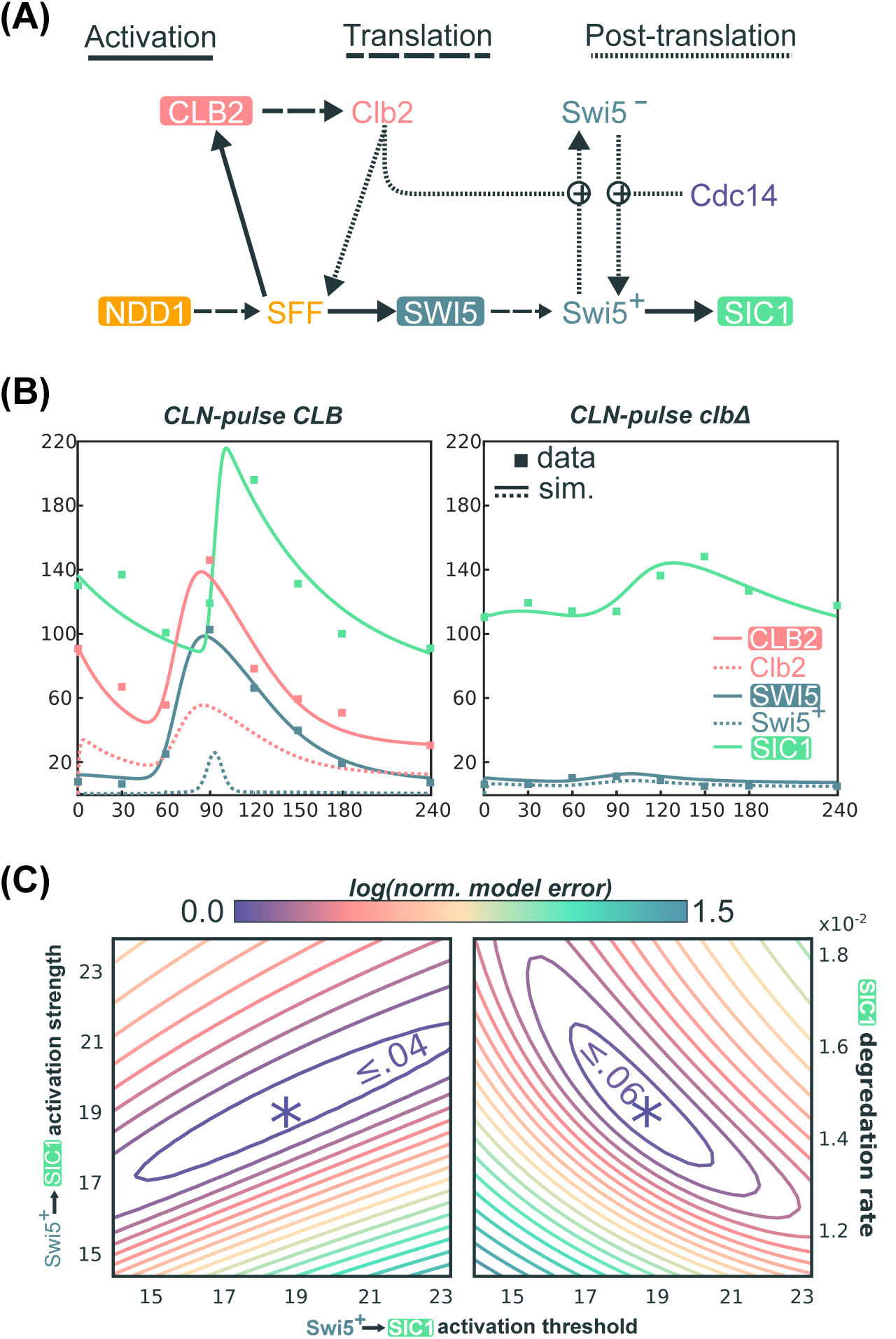
A quantitative model demonstrates the robust activation of *SIC1* by low-amplitude *SWI5* oscillation. (A) Network topology used for quantitative modeling of transcriptional regulation of *SIC1* (see Document S1 for explicit description of equations). (B) Line graphs of selected variables generated by numerical simulation of mathematical model in (A) for a particular choice of parameters, *Θ*\*, along with scatter plots of *CLB2, SWI5,* and *SIC1* levels in the RNA-seq data (in FPKM values) from Rahi et al. (2016). See Document S1 for the parameter values in *Θ*\*. (C) Contour plots of the logarithm of the local-minimum-normalized model error over two 2-dimensional regions of parameters space, centered at Θ*. In particular we plot log(𝓛(Θ)/𝓛(Θ*)), where 𝓛(Θ) is the objective function defined in Document S1, as we independently vary several parameters in a neighborhood of Θ*. See also Figure S4.

To test this hypothesis quantitatively, we constructed a minimal ODE model based only on well-established regulatory interactions (Figure 5A, Materials and Methods; Document S1) (Chen et al., 2004; Kraikivski et al., 2015). Specifically, we aimed to determine whether this model could explain the observation that the roughly 10-fold reduction in *SWI5* peak expression in the *clb*Δ cells does not correspondingly reduce *SIC1* expression. Remarkably, after parameter optimization, this simple model is capable of generating dynamical behaviors of *SWI5* and *SIC1* that closely match the experimental data produced by Rahi et al. (2016) (Materials and Methods and Document S1). For the *CLN-pulse CLB* control cells (Figure 5B, left), the simulations indeed recapitulate the transient burst of *SIC1* expression in late mitosis due to the opposing regulations of Swi5 by Clb2 and Cdc14. Using the same parameters to simulate the *CLN-pulse clb*Δ mutant cells (Figure 5B, right), the model also successfully recapitulates the intermediate activation of *SIC1* by the low-amplitude oscillation of *SWI5.* These dynamical behaviors are achievable by a wide variety of parameter choices, including a large range of activation thresholds for Swi5 activation of *SIC1* (Figures 5C, S4A, and S4B). Finally, similar fits to data can also be observed elsewhere in parameter space (Figure 5C), further supporting the biological plausibility of a model in which substantially reduced levels of Swi5 could still regulate *SIC1* oscillations in *clb*Δ mutants.

While the above model and parameters may not fully represent the physiology of budding yeast cells, these results clearly demonstrate that in the context of a network, a 10-fold drop in expression of a regulator does not necessarily cripple its ability to regulate downstream targets. Given these findings, it is possible that a pulse of transcription is indeed passed through the TF network (Figure 4A). Thus, the parsimonious explanation for the *SIC1* oscillations during CDK-APC/C arrests observed by Rahi et al. (2016)is that they are produced by a TF network that can function in the absence of oscillating CDK activity (Figure 1D).

### The cell-cycle transcriptional program in mitotically arrested cells

The semi-autonomous ability of the TF network to trigger phase-specific transcription has also been tested by a complimentary set of experiments where mitotic cyclins were maintained at high levels (Bristow et al., 2014; Rahi et al., 2016). It has been observed by microarray that transcriptional oscillations continue in the *cdc20*Δ mutant cells arrested in metaphase after *GALL-CDC20* shut-off (Bristow et al., 2014), with a nearly identical period as those observed in the wild-type cells. Nonetheless, it was suggested in a recent study that the transcriptional oscillations observed by Bristow et al. (2014)resulted from cells leaking through the arrest (Rahi et al., 2016). Although several lines of evidence argued against this possibility (see Discussion), we sought to determine whether transcriptional oscillations could be reproducibly observed in other mitotic arrests, such as the anaphase/telophase arrests whose transcriptomic dynamics have not been investigated. To this end, we characterized the global transcript dynamics in the *cdc14-3* and *cdc15-2* mutant cells, which are temperature-sensitive mutants defective in mitotic exit.

In budding yeast, Cdc14 phosphatase is a CDK-counteracting phosphatase and the key effector for anaphase progression and mitotic exit (Amon, 2008; Visintin et al., 1998; Weiss, 2012). The release of Cdc14 from its sequestration in the nucleolus is promoted by mitotic exit pathways upon anaphase entry (Figure 6A). Particularly, APC-Cdc20 initiates the FEAR pathway to trigger the early-anaphase Cdc14 release into the nucleus (Shirayama et al., 1999; Stegmeier et al., 2002; Sullivan and Uhlmann, 2003), while Cdc15 is a component in the MEN pathway that promotes the late-anaphase Cdc14 release into the cytosol (Shou et al., 1999; Visintin et al., 1999). Early G1 cells of the *cdc14-3* and *cdc15-2* mutants were obtained by α- factor arrest at permissive temperature (25°C) and then released into YEP-dextrose medium at restrictive temperature (37°C). Time-series samples were taken and subjected to microarray analysis (Figure 6). The mitotic arrests of the bulk of populations were confirmed by budding indices, DNA content, and the absence of nuclear division (Figure S5).

First, we confirmed that the transcript behaviors of cell-cycle genes were consistent with well-established regulations by Clb-CDKs (Figures 6A and 6B). For example, the SBF-regulated gene (*CLN2*) was indeed expressed at constitutively low level during three different mitotic arrests (Figure 6B), indicating the inhibition of SBF by Clb2-CDK (Amon et al., 1993; Koch et al., 1996). The SFF-regulated gene (*CLB2*) was also impaired in its transcriptional down-regulation, likely due to constitutive Clb-CDK activity. Interestingly, the Swi5-regulated gene (*SIC1*) was weakly expressed in both *cdc20*Δ and *cdc14-3* mutants but was robustly activated in the *cdc15- 2* mutant cells (Figure 6B), suggesting that nuclear Cdc14 can partially activate Swi5 to trigger M-G1 transcription. Importantly, we observed strong re-initiation of the MBF-regulated gene (*POL1*) during these mitotic arrests (Figures 6B and 6C), consistent with the previous observations in the *cdc20*Δ mutant cells (Bristow et al., 2014). Moreover, among the 881 genes shown in Figure 2C, we observed a second transcriptional pulse for a significant proportion of early-cell-cycle genes in the *cdc14-3* mutant cells arrested in anaphase (Figure 6C). Less coherent oscillations of global transcript levels were also observed in the *cdc15-2* mutant cells, suggesting additional roles of nuclear Cdc14 in modulating the dynamics of the TF network.

**Figure 6.**
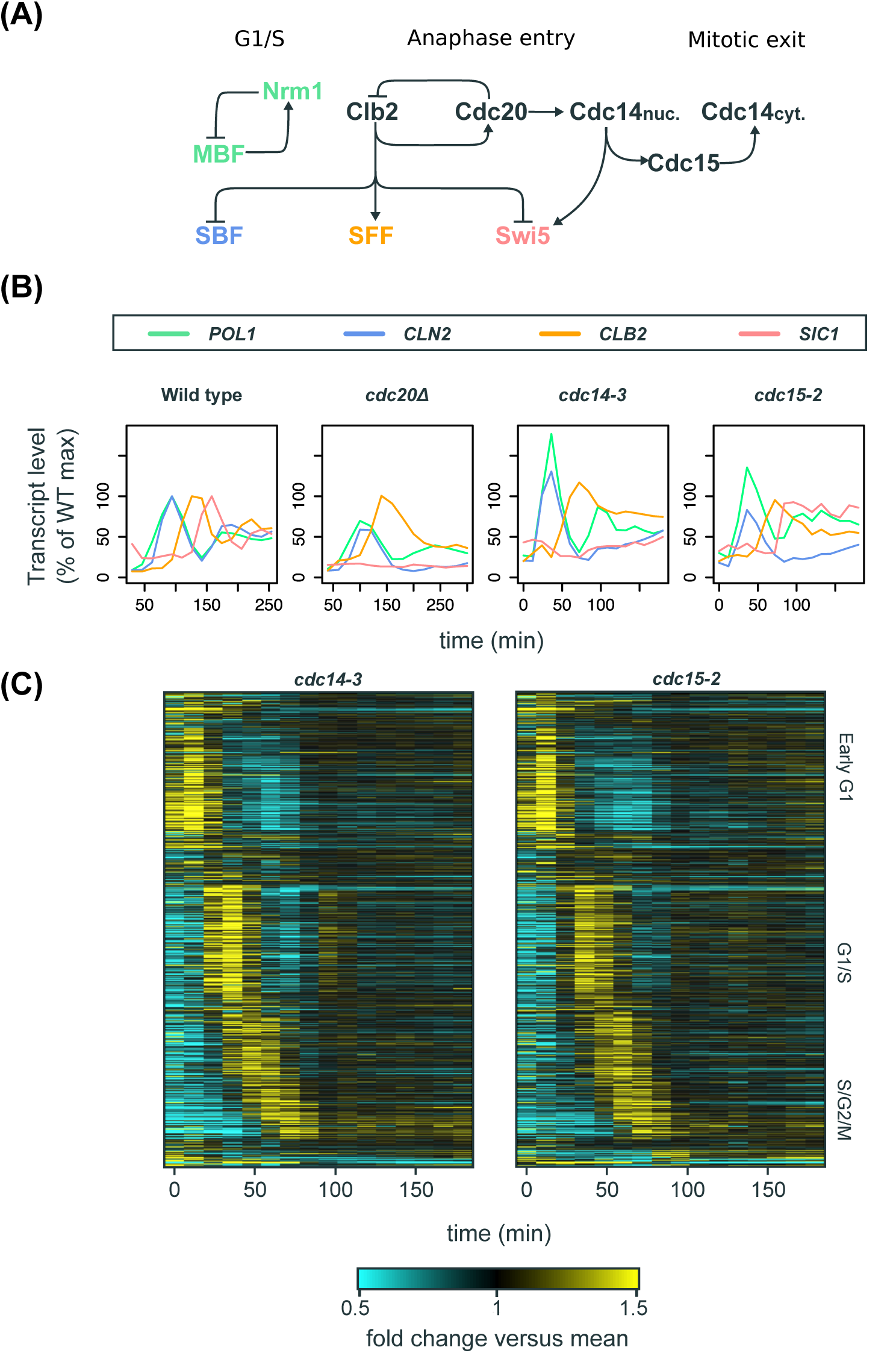
The cell-cycle transcriptional program in mutants defective in mitotic exit. (A) Simplified diagram of part of the CDK-APC/C network (black) controlling progression through mitosis and their established input into network TFs (colored). (B) Line graphs showing transcript dynamics of canonical targets of network TFs in indicated strains. In all experiments, early G1 cells were released for time-series gene expression profiling by microarray. Transcript levels are depicted as percentage of maximal level in corresponding wild-type controls at the same temperature from previous studies (Orlando et al., 2008; Simmons Kovacs et al., 2012). Results of the wild-type control from Orlando et al. (2008)are shown. (C) Heat maps showing the cell-cycle genes shown in Figure 2C in the same order in the *cdc14-3* and *cdc15-2* mutant cells. Early G1 cells were released at restrictive temperature for microarray analysis. Transcript levels are depicted as fold change versus mean in individual dataset. See also Figure S5.

In summary, these data demonstrate again that a large subset of the cell-cycle transcriptional program continues to oscillate in a variety of mutant cells arrested with constitutive Clb-CDK activity, arguing against the model proposed by Rahi et al. (2016) in which oscillations of the CDK-APC/C network predominantly controls global cell-cycle transcription.

## DISCUSSION

Determining how the cell-cycle transcriptional program is generated is important for understanding principles of somatic cell-cycle control. Multiple studies have sought to address this question by monitoring transcript dynamics during a variety of CDK-APC/C arrests (Bristow et al., 2014; Orlando et al., 2008; Rahi et al., 2016; Simmons Kovacs et al., 2012), and two distinct models have been proposed for the global control of cell-cycle transcription, centered on either a network of biochemical CDK-APC/C interactions (Figure 1A) (Rahi et al., 2016) or on a TF network coupled with CDK activities (Figure 1C) (Bristow et al., 2014; Orlando et al., 2008; Simon et al., 2001). We now propose a model of a highly interconnected network (Figures 1D and 7A).

Despite important differences in experimental design, both the Haase and Cross laboratories have performed time-series transcriptome analyses on budding yeast cells deleted for all S-phase and M-phase cyclins (Clb1-6). In the recent study proposing the CDK-APC/C models, Rahi et al. (2016) reported that only three genes were periodically transcribed in the *clb*Δ mutant cells, in contrast to hundreds of periodic genes in the mutant cells from Orlando et al. (2008). Two differences in the analytical approaches likely accounted for the different conclusions drawn in the previous two studies. First, the analysis of Rahi et al. (2016)was restricted to a smaller gene set (91 genes) as compared to a much larger number of periodic genes reported by previous studies (Figure 2B). Second, scores of periodicity test were only computed for clusters of genes by Rahi et al. (2016) rather than for individual genes. Here we found that the only three genes (*SIC1/CDC6/CYK3*) claimed to be oscillating by Rahi et al. (2016)in the *clb*Δ mutant fall to the bottom of the rank-ordered list of periodic genes produced by two distinct periodicity-ranking algorithms (de Lichtenberg et al., 2005; Deckard et al., 2013; Lomb, 1976; Scargle D, 1982), suggesting that one of the major conclusions drawn by Rahi et al. (2016) was internally inconsistent with the RNA-seq data.

By directly comparing the transcriptomic dynamics of an expanded set of 881 genes, we demonstrate that, as observed previously by Orlando et al. (2008), cells lacking B-cyclin genes exhibit a very similar set of dynamics to those with their full complement of B-cyclin genes (Figure 2C). Moreover, the global transcript dynamics are also remarkably similar in the *clb*Δ mutant cells from Orlando et al. (2008) and Rahi et al. (2016) before the *CLN2* shut-off, indicating that residual mitotic cyclin was not responsible for driving transcriptional oscillations as hypothesized by Rahi et al. (2016).

While the dynamics of the first cycle of global cell-cycle transcription look strikingly similar in the *clb*Δ mutant cells from the two studies (Figure 2C), a closer look reveals several differences in canonical gene clusters. For the SBF/MBF cluster, most genes in the cluster did not exhibit a strong second peak of expression in the mutant cells from Rahi et al. (2016), whereas a robust second peak was observed for many SBF/MBF-regulated genes in the mutant cells from Orlando et al. (2008) (Figure 2C and Figure 3). However, the two *clb*Δ mutant strains differed in their expression of G1 cyclins. In the *clb*Δ mutant cells from Rahi et al. (2016), *CLN2* expression was shut-off 90 minutes into the experiment, while *CLN1/2* expression remained high in the *clb*Δ mutant cells from Orlando et al. (2008) (Figure 2A). It is well established that the positive feedback loop mediated by Cln1/2-CDKs removes the transcriptional corepressor Whi5 from SBF complex to promote G1/S transcription (Costanzo et al., 2004; de Bruin et al., 2004; Skotheim et al., 2008). Consistently, the temperature-sensitive *cdc28-4/cdk1* mutant cells only trigger a fraction of the cell-cycle transcriptional program at low amplitude during the G1 arrest (Simmons Kovacs et al., 2012). Thus, a substantial reduction in the amplitude of G1/S transcription after *CLN2* shut-off as observed by Rahi et al. (2016)is fully consistent with these previous findings, and this observation does not rule out the possibility that the TF network can continue to produce phase-specific transcription in the presence of constitutive CDK activity (Figure 1D).

An additional difference in the data derived from the two studies was a greater drop in transcript levels for *CLB2* cluster genes in the *clb*Δ mutant from Rahi et al. (2016). The reduction in transcript levels for these SFF-regulated genes stems from the loss of positive feedback from Clb2-CDK to SFF. The less severe drop observed in the *clb*Δ mutant from Orlando et al. (2008)could potentially result from some residual Clb activity. Regardless, Rahi et al. (2016) asserted that drops resulting in less than 10% of wild-type levels for a TF would render it non-functional. The quantitative models presented here demonstrate the ease with which a 10-fold reduction in the expression level of *SWI5* can continue to drive transcriptional oscillations of *SIC1.* This finding calls into question the validity of the “biological significance” cut-off and supports the TF network models as a plausible explanation for the *SIC1* oscillations observed by Rahi et al. (2016).

Because our ODE formulation represents enzymatic and transcriptional interactions in single cells, the choice to fit the model to RNA-seq data assumes that the populations of cells in these experiments were highly synchronized throughout the time courses. Population modeling of the yeast cell cycle indicates that most of the synchrony loss is due to asymmetric division (Orlando et al., 2009; 2007), and without division in the cyclin mutants, synchrony loss is likely minimal. Given that in the single-cell studies from Rahi et al. (2016), *SIC1pr-YFP* oscillated in the *clb*Δ cells with amplitudes similar to or higher than those in the *CLB* cells but with highly variable peak time, some minor loss of population synchrony could contribute to a reduced *SIC1* peak-to-trough ratio in the RNA-seq data of the *clb*Δ cells. That said, any loss of synchrony should similarly affect the *SWI5* peak-to-trough ratio, so conclusions drawn from the modeling would be largely unaffected by potential loss of synchrony.

Furthermore, when simulating the *clb*Δ mutant using the model presented herein (Figure 5A), it is possible, without changing the expression profile of *SWI5,* to find sets of parameter values yielding even higher amplitudes of *SIC1* oscillations. This is because the activation of *SIC1* can be uncoupled from the accumulation of total Swi5 protein through inhibitory phosphorylation by Clb2-CDK. Thus, a reduction in the activation threshold of *SIC1* by Swi5 will more significantly increase the amplitude of *SIC1* in the model of *clb*Δ mutant cells than in the model of *CLB* cells. It is exactly the strong activation of *SIC1* by Swi5 in concert with the competing actions of Clb2 and Cdc14 on Swi5 that produces the dramatic, switch-like activation of *SIC1* observed in the simple model of *CLB* cells (Figures 5C, S4A, and S4B), while still allowing low levels of Swi5 to drive changes in *SIC1* expression. The ability of the network to produce robust *SIC1* oscillations when *SWI5* transcript levels are reduced by 10-fold suggests that local network motifs make the network robust to perturbations in amplitude, a function previously observed in other network contexts (Acar et al., 2010). Broadly speaking, these results highlight the importance of studying the regulatory interactions in the context of an integrated network.

The CDK-APC/C oscillator model proposed by Rahi et al. (Figure 1A) predicts that if cells were arrested with the CDK oscillator in the “on” state, transcriptional oscillations should stop. In support of this prediction, it was demonstrated in single-cell assays that *CLN2pr-GFP* and *CLB2pr-GFP* did not oscillate in cells arrested with high Clb-CDK activity (Rahi et al., 2016). It was confusing that the authors limited the single-cell analyses to these two genes, as it was shown previously by microarray analysis that neither *CLN2* or *CLB2* transcripts oscillated in Cdc20-depleted cells, despite the fact that many other transcripts continued to oscillate (Bristow et al., 2014).

In support of previous findings, we now demonstrate that substantial transcriptional oscillations persist in mitotic exit mutants where the cell cycle is arrested and Clb2 is maintained constitutively at moderate levels (Figure 6) (Bäumer et al., 2000; Visintin et al., 1998; Yeong et al., 2000). We argue that the transcriptional oscillations observed in these mitotically arrested cells (and the Cdc20-depleted cells from Bristow et al., 2014) did not result from cells leaking through the arrest for the following reasons. First, as indicated by budding indices and the Clb2 level in western blots (Figure S5; Bristow et al., 2014: Figure S1), the bulk of the population remained mitotically arrested throughout the experiments, and thus very few cells leak through the arrest. Moreover, the second cycle of transcription exhibited amplitudes very similar to wild type (Figure 6C; Bristow et al., 2014: Figure 3), which is not consistent with a small population of cells leaking through the arrest. Finally, the transcript behaviors of established Clb2-CDK transcriptional targets were indeed impaired during the arrest, including the SBF-regulated genes (inhibited by Clb2; Amon et al., 1993; Koch et al., 1996) and the SFF-regulated genes (up-regulated by Clb2; Pic-Taylor et al., 2004; Reynolds et al., 2003) (Figure 6B; Bristow et al., 2014: Figure 1), consistent with the results of the single-cell assays (Rahi et al., 2016).

Collectively, the data argue against the CDK-APC/C models (Figure 1A) in which periodic input from CDKs is needed to drive periodic cell-cycle transcription. The data are consistent with an integrated model (Figure 1D) in which CDK-APC/C and the TF network function together to drive the cell-cycle transcriptional program during the normal cell cycle (Figure 7A). The high degree of interconnection between network TFs and CDKs argues against models where one autonomous oscillator is entraining another. Such an integrated network would couple transcriptional oscillations with normal cell-cycle progression as well as promote robustness of cell-cycle oscillations to a variety of perturbations.

**Figure 7.**
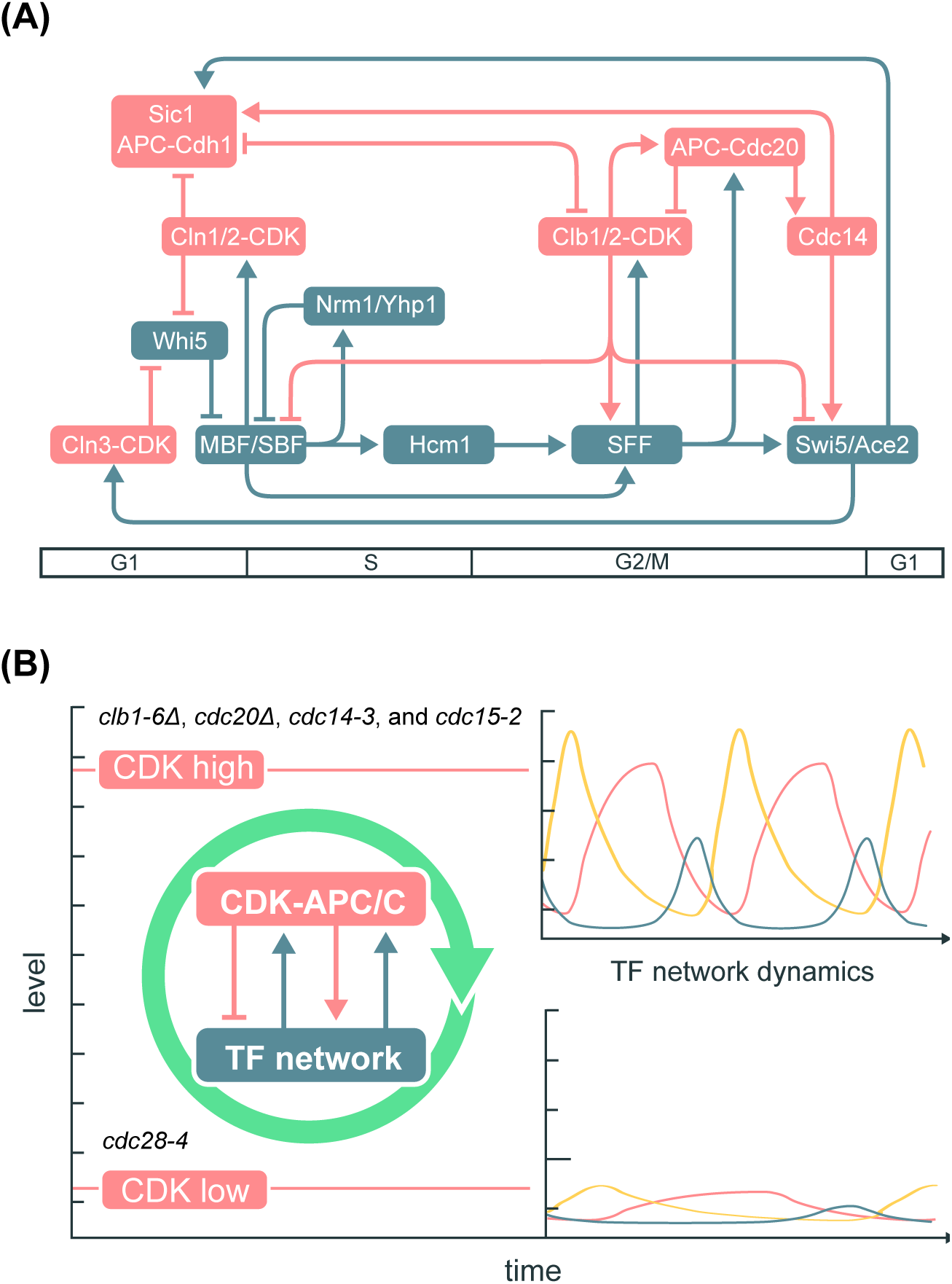
Integrated network model for the control of the cell-cycle transcriptional program in budding yeast. (A) Network diagram incorporating components of the CDK-APC/C model proposed by Rahi et al. (2016) and the TF network model proposed by Orlando et al. (2008). Nodes are ordered horizontally by their approximate time of activation during the cell cycle. (B) Functional outcomes of the cell-cycle-transcriptional program during different CDK-APC/C perturbations with either high or low CDK activities.

Moreover, this integrated model can explain the transcript dynamics observed in multiple mutant backgrounds (Figures 2C and 6C). In the CDK-APC/C mutants, low levels of CDK activities are then expected to impair the capability of the TF network to generate the cell-cycle transcriptional program, while constitutively high CDK activities can promote the TF network to generate global cell-cycle transcription even without oscillating CDK activities (Figure 7B). The *CLN-pulse clb*Δ experiments from Rahi et al. (2016)can then be viewed as a hybrid of high- and low-CDK arrests sequentially, and thus only one robust cycle of transcription could be observed. An interesting question to ask is whether the impaired dynamics of the TF network in low-CDK conditions could be genetically restored, such as by deleting the transcriptional corepressor (Whi5). Analogously, the lethality of *cln1 cln2 cln3* triple mutant can be rescued by the *sic1* mutation that restores B-cyclin-CDK activities (Schneider et al., 1996; Tyers, 1996).

We have proposed that the ancestral oscillatory mechanism for the cell cycle was a TF network (Simmons Kovacs et al., 2008), while CDKs have been proposed to arise in evolution well after mechanisms of cell division had been established (Krylov et al., 2003). In modern eukaryotes, CDKs and APC/C undoubtedly provide important feedback regulations onto the TF network to modulate the cell-cycle transcriptional program (Figure 6). Dissecting and establishing the molecular mechanisms that couple the oscillations of CDK-APC/C and the TF network will be imperative for moving toward an integrated model of the eukaryotic cell cycle.

### Materials and Methods

Requests of further information may be directed to the corresponding author Steven B. Haase.

#### Processing and analyses of RNA-seq data

Raw RNA-Sequencing data from Rahi et al. (2016)were downloaded from the SRA database (http://www.ncbi.nlm.nih.gov/sra/?term=SRP073907). FASTQ files were aligned using STAR (Dobin et al., 2013). The S. *cerevisiae* S288C reference genome (Ensembl build R64-1-1) was downloaded from Illumina iGenomes on March 2, 2016 (https://support.illumina.com/sequencing/sequencing_software/igenome.html). Aligned reads were assembled into transcripts, quantified, and normalized using Cufflinks2 (Trapnell et al., 2013). Samples from all time-series experiments were normalized together using the CuffNorm feature. Replicate time-series data were available on SRA, but not discussed in detail by Rahi et al. (2016). Therefore, we relied on the SRA annotation to organize them as summarized in Table S1.

The normalized FPKM gene expression outputs (“genes.fpkm_table”) were used in the analyses presented. To avoid fractional and zero values, 1 was added to every FPKM value in each dataset. Fractions and zeros were found to interfere with the de Lichtenberg periodicity algorithm, which involves log-transformation of data points (data not shown).

Four time-series datasets corresponding to the *CLB* control and clbΔ mutant experiments (two replicates each) described in Rahi et al (2016) were run through two periodicity-ranking algorithms: Lomb-Scargle (LS) and de Lichtenberg (DL) (de Lichtenberg et al., 2005; Lomb, 1976; Scargle D, 1982). Each algorithm was implemented as described previously (Deckard et al., 2013). For all time-series data, we first tested a large range of periods from 50–130 minutes as previously described (Rahi et al., 2016). We examined the top-scoring genes in the LS output for each replicate dataset. At a p-value cutoff of 0.2, about 600–900 genes were reported as periodic by LS. This numerical p-value cutoff is arbitrary, but the size of the output periodic gene lists matches previous literature (Bristow et al., 2014; Cho et al., 1998; de Lichtenberg et al., 2005; Eser et al., 2014; Granovskaia et al., 2010; Kelliher et al., 2016; Orlando et al., 2008; Pramila et al., 2006; Spellman et al., 1998). For those top periodic genes, we then examined their period length reported by LS in the following table.

| Experiment | Number_genes_LS<0.2 | Average_period_LS<0.2 | Stdev_period_LS<0.2 |
| --- | --- | --- | --- |
| *CLN*-pulse *CLB* rep1 | 615 | 95.4 | 26.4 |
| *CLN*-pulse *CLB* rep2 | 617 | 96.4 | 26.8 |
| *CLN*-pulse *clbΔ* rep1 | 971 | 99.4 | 25.8 |
| *CLN*-pulse *clbΔ* rep2 | 909 | 100.1 | 25.1 |

These results indicated that the dominant periods in the “most periodic” gene set are 100 ± 30 minutes. This new range eliminates the original lower bound of 50 minutes used by Rahi et al (2016). However, the period length of cycling wild-type cells in rich media is longer than 50 minutes. Although Rahi et al (2016)did not report bud emergence timing for these experiments, the cells were cultured in synthetic media but not rich media. Therefore, an average period length of 100 minutes for *CLN-pulse clb*Δ and CLN-pulse *CLB* cells seemed to be a reasonable estimate. Thus, we ran DL at the average period length of 100 minutes, and re-ran LS at a range of 70–130 minute periods. We used the second set of DL and LS results to search for and rank periodic genes in each experiment from Rahi et al. (2016) (Table 1).

#### Yeast strains and cell culture synchronization

The *cdc14-3* and *cdc15-2* strains are derivatives of S. *cerevisiae* BF264-15D (*ade1 his2 leu2-3,112 trp1-1a*). Strain A1268 (W303 *cdc14-3 PDS1-HA-LEU2::pds1*) was provided by Angelika Amon (Visintin et al., 1998) and outcrossed with BF264-15D for 5 times. Yeast cultures were grown in standard YEP medium (1% yeast extract, 2% peptone, 0.012% adenine, 0.006% uracil supplemented with 2% sugar). For synchronization by α-factor, cultures were grown in YEP-galactose medium at 25°C and incubated with 50 ng/ml α-factor for 140–180 minutes. Synchronized cultures were then resuspended in YEP-dextrose medium at 37°C. Aliquots were taken at each time point and subsequently assayed by microarray or western blots.

#### RNA extraction and microarray assay

Total RNA was isolated by standard acid phenol protocol and cleaned up by RNA Clean and Concentrator (Zymo Research) if necessary. Samples were submitted to Duke Microarray Facility for labeling, hybridization, and image collection. mRNA was amplified and labeled by Ambion MessageAmp Premier kit (Ambion Biosystems) and hybridized to Yeast Genome 2.0 Array (Affymetrix).

#### Microscopy

Cells were fixed in 2% paraformaldehyde for 5 minutes at room temperature, washed with PBS, and then resuspended in 30% glycerol for mounting on glass slides. All imaging was performed on Zeiss Axio Observer.

#### Flow cytometry

Cells were prepared for flow cytometric analysis using SYTOX Green staining as described (Haase and Reed, 2001). Graphs were generated using the FlowViz package in Bioconductor in R.

#### Normalization of microarray data

Previously published datasets used in this study are GEO: GSE8799, GEO: GSE32974, and GEO: GSE49650. All CEL files analyzed in this study were normalized together using the dChip method from the Affy package in Bioconductor as described previously (Bristow et al., 2014).

#### Quantitative modeling of *SIC1* activation

Justifications and full methodology of the mathematical modeling can be found in Document S1.

## ACKNOWLEDGEMENTS

We thank Daniel J. Lew and Adam R. Leman for helpful discussions and critical readings of the manuscript.

